# Mecp2 deficiency impairs microscale cortical network topology and dynamics in a Rett syndrome mouse model

**DOI:** 10.64898/2025.12.17.694950

**Authors:** Alexander W. E. Dunn, Timothy P. H. Sit, Rachael C. Feord, Asma Soltani, Yin Yuan, Richard Turner, Ines Loureiro, Stephen J. Eglen, Ole Paulsen, Susanna B. Mierau

## Abstract

Rett syndrome is a debilitating neurodevelopmental disorder with cerebral processing impairments caused by *MECP2* loss-of-function mutations. Mecp2-deficient mouse models reveal disruptions of microscale cortical circuits. Yet how cellular-scale information processing is altered in Mecp2-deficient microscale functional networks is unknown. We investigated the development of functional connectivity, network topology, and dynamics in microelectrode array (MEA) recordings of primary cortical cultures from Mecp2-deficient and wild-type mice. Mecp2-deficient cortical networks developed more slowly and showed decreased functional connectivity compared to wild-type, leading to smaller network size, density, and strength of connectivity. Altered network topological features in Mecp2-deficient microscale circuits predicted decreased efficiency and information-sharing capacity. This reveals developmental deficits in microscale functional networks, which may in turn underlie the cortical decline and severe cognitive disability in Rett syndrome. These findings also offer circuit-level targets and an in-vitro approach for evaluating new therapeutic products for restoring microscale network function.

## Introduction

Rett syndrome is a neurological disorder in which children, primarily girls, have severely impaired cerebral processing leading to lifelong deficits in cognitive, language, social, motor, and sensory function [1]. Although typically diagnosed after 6-18 months of age, behavioral differences are already apparent at birth [2]. Loss-of-function mutations in the gene regulator *MECP2* were first identified 25 years ago and account for 95% of Rett syndrome [3]. At the whole-brain or cortical-region-levels, impairments have been identified in cortical processing, using visual evoked potentials [4], and network architecture [5]. However, how MeCP2 deficiency alters the development of the brain networks that support higher cognitive functioning is yet unknown, and there is no treatment that can stop the decline or reverse the deficits.

Mecp2-deficient mice recapitulate many behavioral features of the human disorder [6] and display synaptic deficits that precede the behavioral decline [7–15]. In particular, loss of Mecp2 disrupts the timing of cortical excitatory and inhibitory cell-type maturation, delaying maturation in excitatory neurons and accelerating maturation in parvalbumin-positive (PV) inhibitory neurons [15]. This disruption in synaptic maturation would likely impact the formation of functional connectivity in the developing brain. The organization of neurons into functional microscale networks is key to the computational and learning capabilities of the brain [16]. These patterns of network topology are seen across spatial scales in the brain and determine the efficiency of information processing [17]. Whole-brain to regional-level alterations in the degree of functional connectivity in the cortex of Mecp2-deficient mice, visualized using optical imaging, support network-level defects [18]. Anatomical tracing also confirms impairment in long-range connectivity in Mecp2-deficient hippocampal circuits [19]. These differences in functional and structural connectivity at larger spatial skills likely arise, at least in part, from altered maturation of microscale networks. Microelectrode array (MEA) recordings reveal that primary murine cortical cultured networks can process complex spatiotemporal information [20]. Thus, identifying how Mecp2 deficiency alters the development of microscale network topology and dynamics in vitro could reveal mechanistic insights and novel therapeutic targets for Rett syndrome.

To address this gap, we compared the development of functional connectivity and network topology in developing primary cortical cultures from mice hemizygous (KO, male) or heterozygous (HET, female) for a loss-of-function deletion of exon 3 and 4 in *Mecp2* on the X chromosome and their wild-type (WT) littermates. We found that Mecp2 deficiency reduces and delays the development of spontaneous activity in developing cortical microcircuits affecting firing rates, network burst rates, and functional connectivity in both the KO (all cells lack Mecp2), and HET (mosaic for Mecp2 expression due to X-inactivation) cultures. Mecp2 deficiency affects the network topology, including nodal- and recording-level network features. These topological features predict reduced information processing capacity in Mecp2-deficient microscale cortical networks, which we confirmed by identifying subnetworks—based on their patterns of activity—using dimensionality reduction techniques.

## Results

### Mecp2 deficiency alters developmental trajectory of spontaneous activity

We first compared the development of the number of active electrodes and firing rates in the cortical cultures over days-in-vitro (DIV) 14-35 (**Figure 1, Table S1**). Notably, due to X-inactivation, while all cells in the Mecp2-WT cultures expressed Mecp2, there is mosaic expression of Mecp2 in the HET cultures, and no cells expressed Mecp2 in the KO cortical networks (**Figure 1A**). Although the number of active electrodes and the median firing rates increased over development in all three genotypes, the Het and KO cultures had fewer active electrodes and lower median firing rates at DIV 14-35 (p<0.05, nparLD; **Figure 1B-D**). This suggests that loss, or partial loss, of Mecp2 function delays and reduces the developmental delay in spontaneous neuronal activity. This effect may involve non-cell-autonomous effects, as there were similar effects on number of active electrodes, and firing rates at the later time points, in the HET cultures, which are mosaic for Mecp2 expression, and the KO cultures, which lack Mecp2.

**Figure 1.**
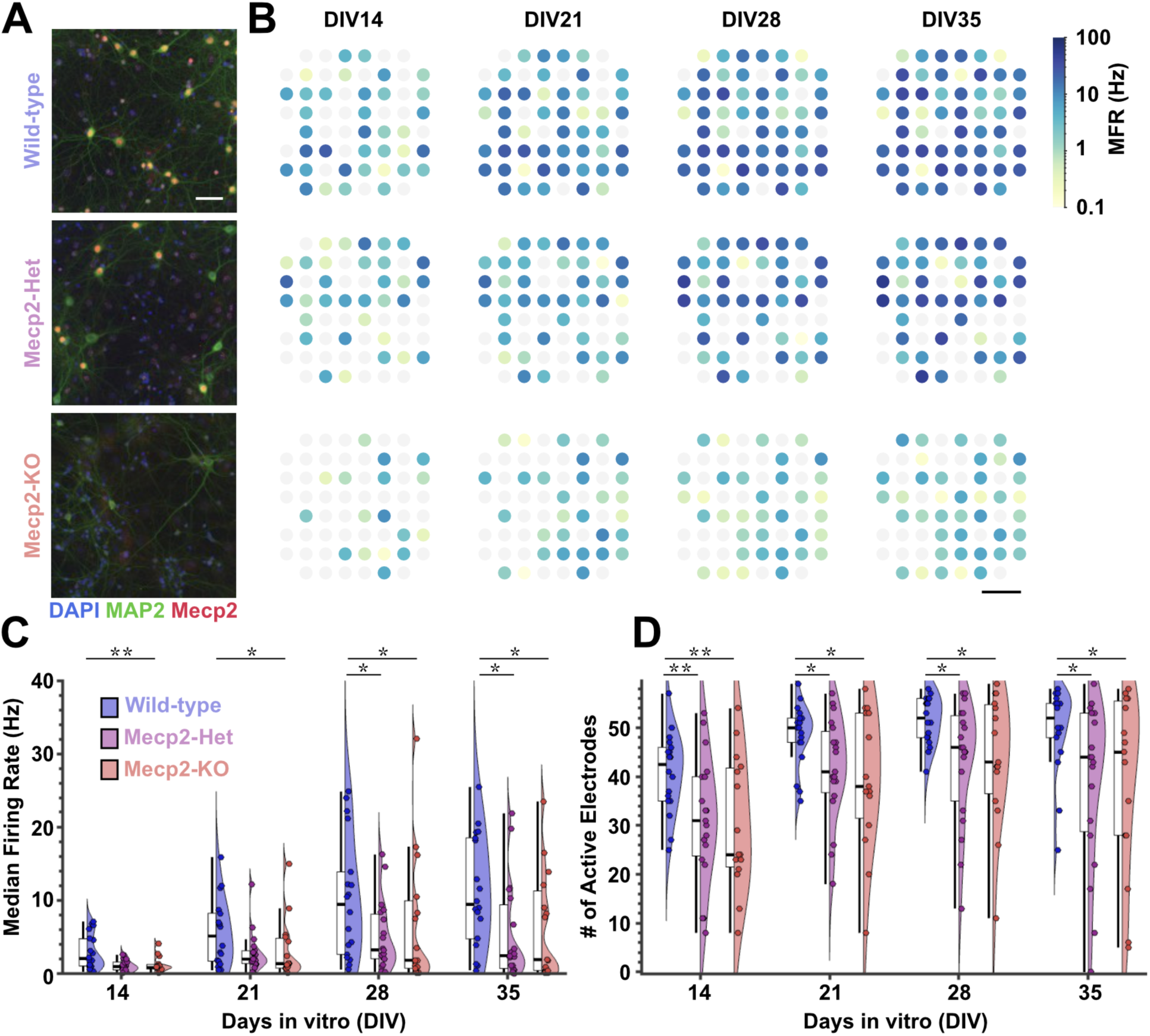
Mecp2 deficiency alters the development of spontaneous activity in murine cortical cultures, see also Table S1. **A.** Representative images of Mecp2-wildtype (WT), - heterozygous (Het) and -hemizygous (KO) cortical cultures at DIV15 with immunostaining for Mecp2 (red), MAP2 (green), and DAPI (blue) reveal Mecp2+ (yellow) and Mecp2- (green) cells in the HET culture. Scale bar, 40 μm. **B.** Representative mean firing rate heat maps in the spatial arrangement of the microelectrode array (MEA) at days-in-vitro (DIV) 14 to 35 from Mecp2-WT (top row), -Het (middle row), and -KO (bottom row) cortical cultures. Scale bar, 400 μm. Node color shows the mean firing rate (MFR; action potentials per second) for each electrode in the 10-minute MEA recordings. **C-D.** Box plots show the median (horizontal line), interquartile range (box), and range, excluding outliers (vertical lines), with scatter plots (colored circles for each culture) and density curves for the median firing rate for each culture (**C**) and number of active electrodes (**D**). Statistical significance was assessed using nparLD, followed by Dunn’s Test for post hoc comparisons. * p≤0.05, ** p≤0.01.

We next compared single-electrode and network bursting in the Mecp2 WT, Het, and KO cortical cultures (**Figure 2, Table S2**). Neurons in culture develop a pattern of firing in which action potentials occur in bursts, with shorter interspike intervals (ISI) within bursts and longer ISI between bursts (**Figure 2A-C**). We first compared bursting that occurred in single electrodes (**Figure 2D-E**). The percent of electrodes showing bursting activity increased with development in all three genotypes (**Figure 2D**). The WT cortical networks showed a higher percentage of electrodes bursting than the HET and KO cultures at DIV14-35. The mean burst rate in individual electrodes also increased with development (**Figure 2E**). Interestingly, in the HET and KO cultures, the neurons that did show single-electrode bursting achieved the same mean rate as in WT.

**Figure 2.**
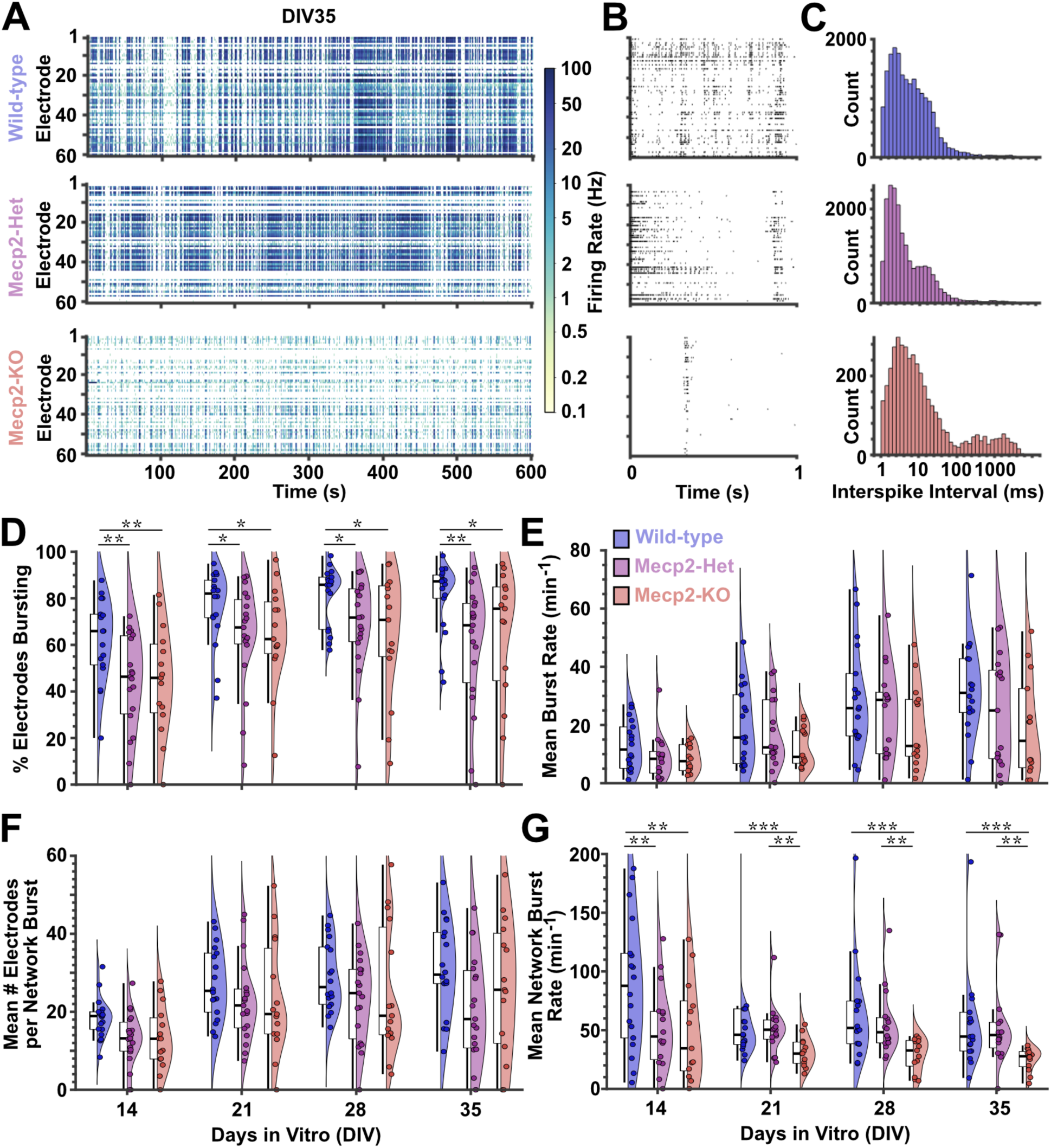
Mecp2 deficiency alters the development of single-electrode and network bursting in murine cortical cultures, see also Table S2. A. Representative raster plots of spontaneous activity from 10-minute MEA recording from Mecp2-wildtype (WT, top), heterozygous (Het, middle) and hemizygous (KO, bottom) cortical cultures at days-in-vitro (DIV) 35 showing bursting in individual electrodes and network bursts. Color bar shows firing rate (action potentials per second). B. Representative raster plots of 1-second periods from each recording illustrating the different patterning of bursting activity in the three genotypes. Each thin vertical line represents an action potential and each row represents an electrode in the same recording. C. Histograms of interspike intervals (ISI) in milliseconds (ms) for each of the recordings in A. D-G. Box plots show median (horizontal line), interquartile range (box), and range excluding outliers (vertical lines) with scatter plots (colored circles for each culture), and density curves for the mean percent of electrodes bursting (D), mean single-electrode burst rates per minute (E), mean number of electrodes participating in network bursts (F), and network burst rates per minute (G) for DIV 14-35. * p<0.05, ** p≤0.01, *** p≤0.001 (nparLD, Dunn’s Test).

Bursts that occur simultaneously in multiple electrodes are called network spikes or bursts [21] and arise from the increase in functional connectivity within the cortical networks. The raster plots illustrate the differences in patterning of burst activity at the whole-recording (**Figure 2A**) and one-second temporal windows (**Figure 2B**). WT and HET networks show complex bursting that is apparent in the 1-second temporal window and ISI distribution plots (**Figure 2B-C**). In contrast, the KO cultures show smaller network bursts with much less firing in between the network bursts, as illustrated by the ISIs greater than 100 milliseconds. The mean number of electrodes participating in network bursts increased in all three genotypes at DIV14-35 (**Figure 2F**). However, the WT cultures showed higher mean network burst rates than HET cultures at DIV14 and KO cultures at DIV14-35, while the HET cultures showed higher mean network burst rates than KO cultures at DIV21-35 (**Figure 2G**). Thus, loss of Mecp2 altered not only the firing rate, but also the patterning of spontaneous activity in in vitro developing cortical networks. There may be non-cell-autonomous effects of Mecp2 mosaicism on the development of bursting in the HET networks, as the mean network burst rate differed from the WT at DIV14 and KO at DIV21-35.

### Mecp2 deficiency impairs the development of functional connectivity

To determine whether Mecp2 deficiency alters network function in microscale circuits, we first compared the functional connectivity using the spike time tiling coefficient (STTC) [22] and probabilistic thresholding [23]. By determining significant pairwise correlations of neuronal activity detected at the electrodes, we could reliably infer significant functional connections. These networks can be visualized as nodes (activity recorded from each electrode) and edges (significant correlated activity observed between two nodes) in the spatial arrangement of the MEA (**Figure 3A**) and network features compared (**Figure 3B-3F, Table S3**). We found that Mecp2 deficiency impairs the development of functional connectivity in the cultured cortical networks. The HET and KO networks show a reduction in the mean edge weight (strength of connectivity; **Figure 3C**), mean node degree (number of connections per node; **Figure 3D**), network size (number of nodes; **Figure 3E**), and network density (number of connections as a percentage of total possible connections; **Figure 3F**). The developmental increase in network size, density, and edge weights was slower in the Het and KO networks, compared to WT, and was reduced compared to WT at DIV 14-35. This indicates that, in addition to lower firing and burst rates, Mecp2 deficiency also reduces the degree of pairwise correlations in the timing of action potentials between electrodes. Mecp2-deficient networks were smaller (fewer nodes) and had fewer and weaker connections than WT networks.

**Figure 3.**
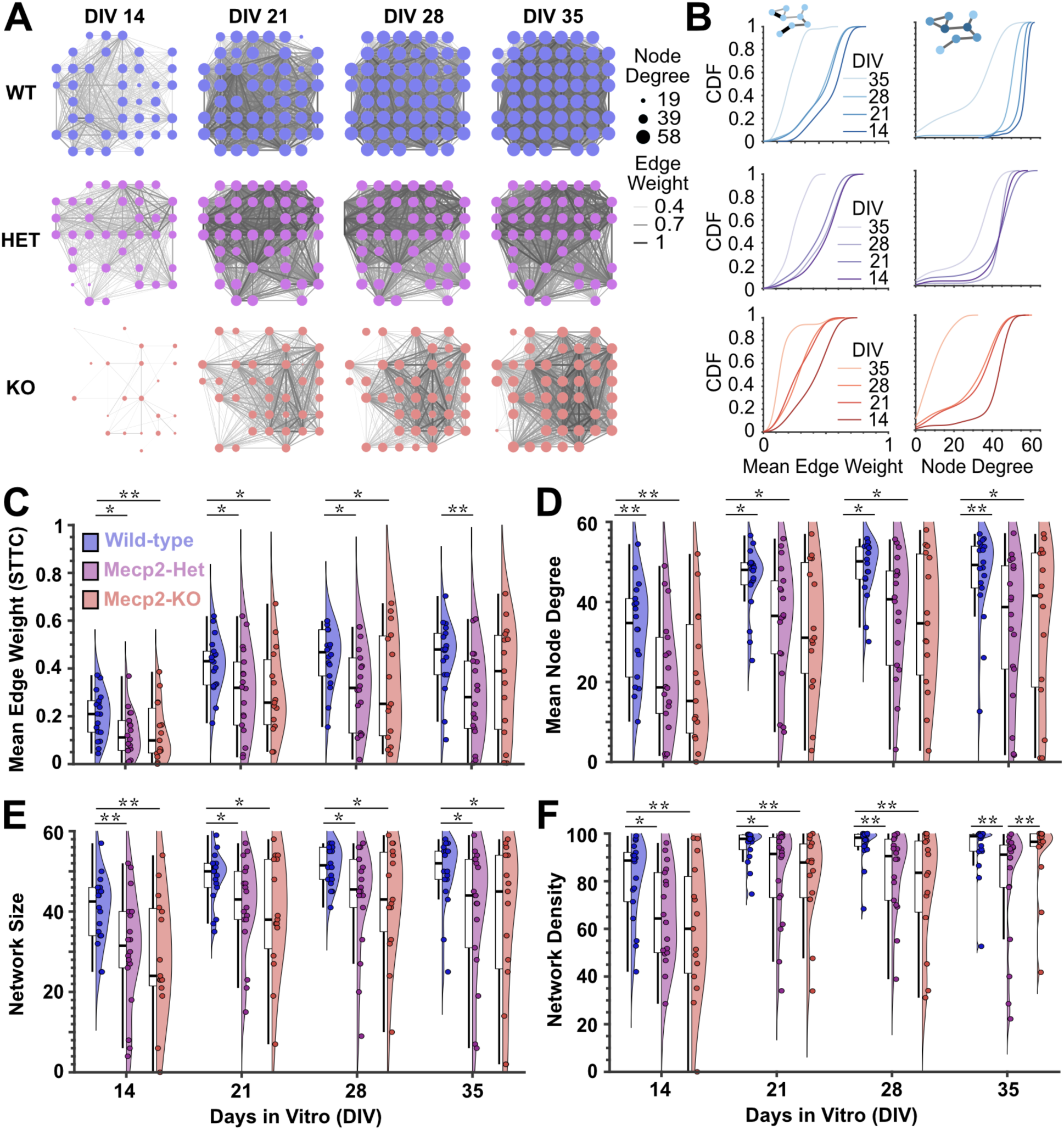
Mecp2 deficiency delays the development and reduces the number and strength of functional connections, see also Table S3. **A.** Representative graphs of networks at DIV 14-35 from *Mecp2*-wildtype (WT,blue), -heterozygous (Het, purple), and -hemizygous (KO, red) cultures. Nodes (circles) are in the spatial arrangement of the MEA. Edge weight (line thickness) indicates the strength of connectivity using the spike time tiling coefficient (STTC). Node size is relative to node degree (number of edges). **B.** Cumulative distribution function (CDF) plots for mean edge weight (left) and node degree (right) shows the developmental shift with age (different line shades) for the cultures in A. Small network schema illustrate edge weight (left, line thickness indicates stronger connection) and node degree (right, darker blue nodes have more connections than lighter blue nodes). **C-F.** Box plots show median (horizontal line), interquartile range (box), and range excluding outliers (vertical lines) with scatter plots (colored circles for each culture), and density curves for the mean edge weight (**C**), mean node degree (**D**), network size (**E**), and network density (**F**) for days-in-vitro (DIV) 14-35. * p≤0.05, ** p≤0.01 (nparLD, Dunn’s Test).

### Mecp2 deficiency alters development of rich-club hubs and small-world topology

To determine how Mecp2 deficiency alters the network topology—the organizational patterns or motifs observed in the functional connectivity—we analyzed multiple graph theoretical metrics (**Figure 4, Table S4**). The distribution of functional connections (edges) between nodes is not uniform. There are clusters of nodes, for example, that form subcommunities within the overall network activity and within these subcommunities there are nodes that serve as hubs, because they have greater connectivity to the other nodes. Hub nodes from different subcommunities can also cluster together to form “rich clubs.” Rich clubs increase in WT networks between DIV 14 to 21 and maintain their dense connections through DIV 35. In contrast, rich clubs in HET and KO networks increased more slowly (**Figure 4A**). The distribution of the clustering coefficient and the betweenness centrality (proportion of shortest paths between any two nodes that goes through a given node) also showed developmental shifts (**Figure 4B**). The number of rich club nodes increased with age and was higher in the WT than the HET and KO at DIV 14-35 (**Figure 4C**). The betweenness centrality, normalized to surrogate graphs, revealed differences between the WT and HET at DIV 21-28 (**Figure 4D**). The rich club coefficient (normalized to surrogate graphs) was lower in the WT than HET or KO at DIV14-28 (**Figure 4E**). The presence of clustered subcommunities and rich-club hubs connecting them can promote more efficient information processing in networks by reducing the energy costs. The WT showed a normalized small-world coefficient approaching 1, which was significantly different from the higher small-world coefficients in the HET and KO networks at DIV 21-28 (**Figure 4F**).

**Figure 4.**
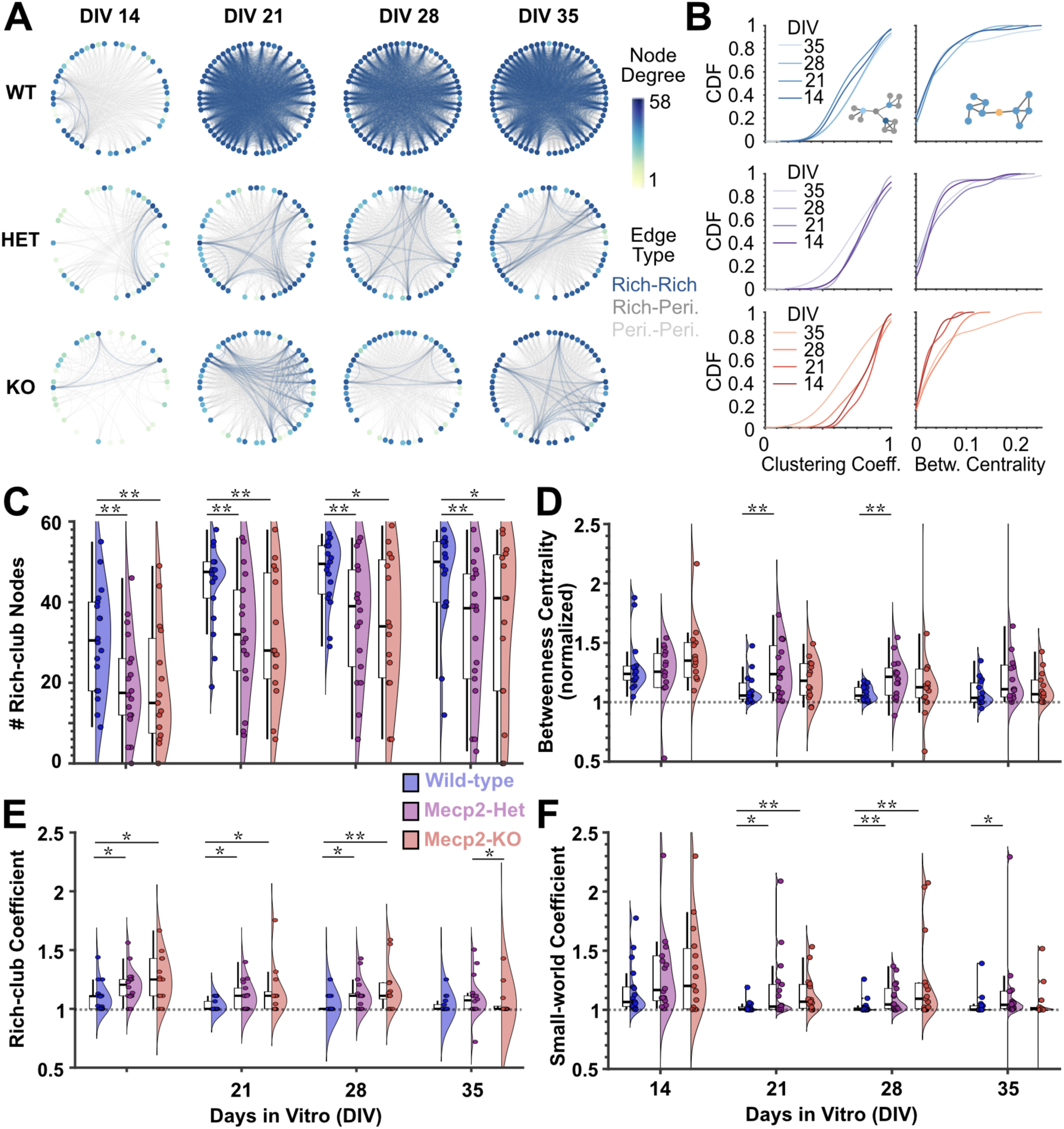
Mecp2 deficiency impairs development of rich-club hubs and small-world topology, see also Table S4. **A.** Representative circular network plots show development of rich-club nodes in which node degree (number of connections per node, color bar) and edge type (color) are shown. The dark blue nodes are highly connected, and the dark blue edges show the rich-club-to-rich-club-nodes connections. The less connected nodes are lighter colors. The connections between rich-club and peripheral (peri.) nodes are in dark grey and between peripheral nodes are in light gray. **B.** Cumulative distribution function (CDF) plots for clustering coefficient (left) and betweenness centrality (right) shows the developmental shift with age (different line shades) for the cultures in A. Small network schema illustrate clustering coefficient and betweenness centrality. **C-E.** Box plots show median (horizontal line), interquartile range (box), and range excluding outliers (vertical lines) with scatter plots (colored circles for each culture), and density curves (right) for number of rich-club nodes (**C**), betweenness centrality (**D**), rich-club coefficient (**E**), and small-worldness coefficient (**F**). Cultures without rich clubs (rich-club coefficient=0) are not included in E (n=1 HET and 1 KO at DIV14, n=2 KO at DIV35). * p≤0.05, ** p≤0.01 (nparLD, Dunn’s Test).

### Mecp2 deficiency impairs the development of network dynamics in cultured cortical microcircuits

The altered trajectories of network development we identified above predict an impaired capacity of the Mecp2-deficient networks to support information processing at the microscale. To test this, we applied a dimensionality reduction approach, non-negative matrix factorization (NMF), to identify the number of patterns of activity (NMF components) detected in the MEA recordings (**Figure 5, Table S5**). For the NMF analysis, the entire spike-time time series was used. In contrast to the graph theoretical analysis performed, the dimensionality reduction did not rely on pairwise functional connectivity. Instead, the method identified patterns of activity observed within the entire MEA recording (**Figure 5A**), including patterns which may be detected in only a subset of the electrodes. To quantify the information-processing capacity of the networks, we determined the number of significant activity patterns by first calculating the mean square root residual (MSRR) for different numbers of NMF components (k=1,2,..,50) for the observed network activity and randomized network activity (created by circular shifts of the electrode time series) and second identifying the number of NMF components at which the MSRR in the observed activity was greater than in the randomized network (**Figure 5B**). We found that Mecp2-deficient (HET and KO) networks support fewer patterns of activity than the wild-type networks as determined by a smaller number of significant NMF components (**Figure 5C**). The mean size of the subnetworks per activity pattern (number of electrodes detecting activity per significant NMF component) increased over development in all three genotypes; however, the KO subnetworks had fewer electrodes than HET and WT at DIV14 on average and the HET and KO subnetworks were smaller on average than the WT at DIV21 and 35. (**Figure 5D**). These results suggest that Mecp2-deficient networks have less information-sharing capacity, which may underlie impairments in cortical processing in Rett syndrome.

**Figure 5.**
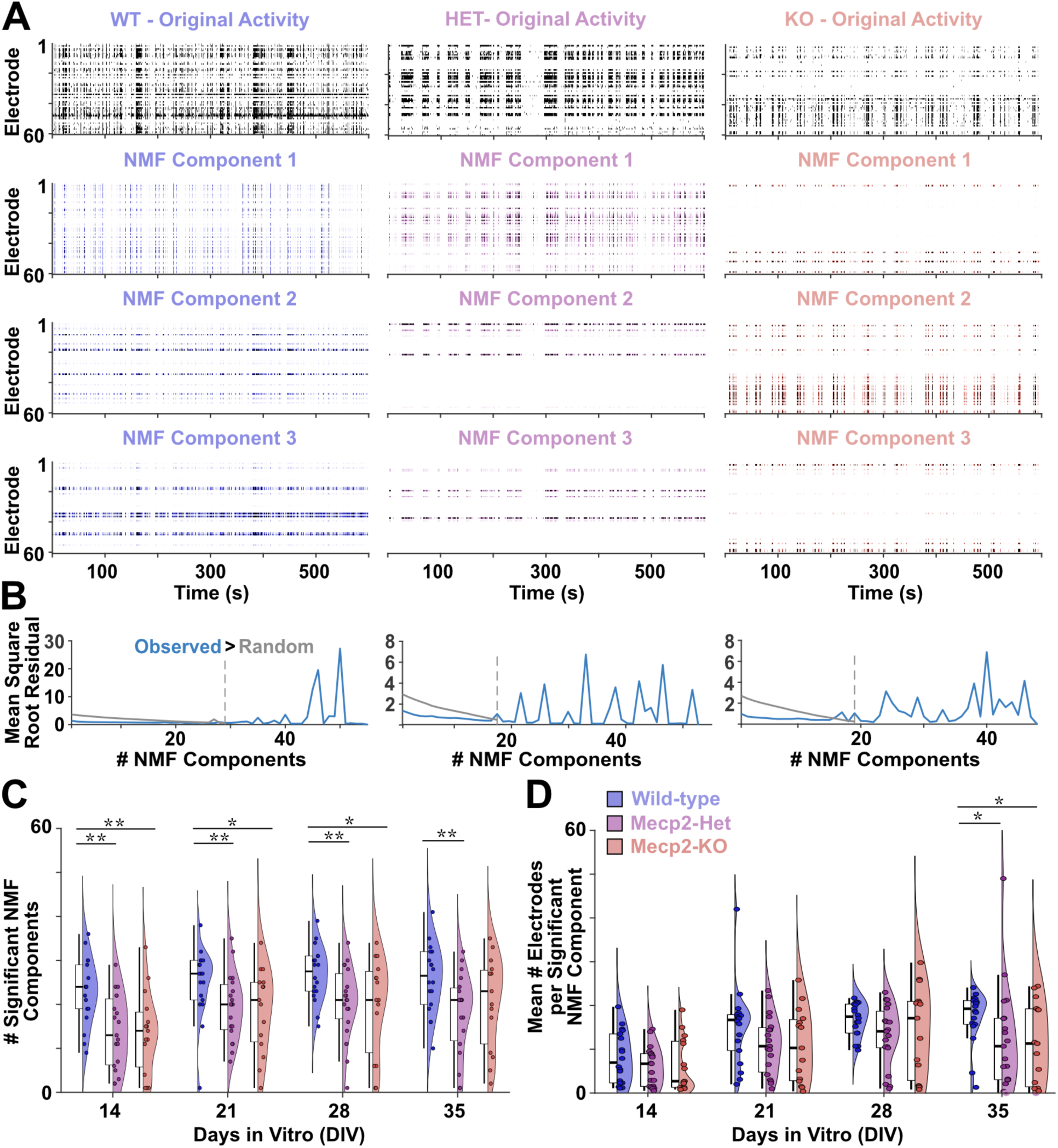
Mecp2 deficiency impairs temporal dynamics in in vitro cortical networks, see also Table S5. **A.** Representative raster plots of the observed spontaneous activity (top row, black) and the top three non-negative matrix factorization (NMF) components (lower 3 rows, color) ranked based on the percent of the variance explained for Mecp2-wildtype (WT, blue), heterozygous (HET, purple), and hemizygous (KO, pink) networks at DIV 35. Each hash represents an action potential; time bins with more than one spike shown in darker shades. In the examples shown, the percent of the variance explained by the top 3 NMF components are 90%, 4%, 1% for WT, 87% 4%, 3% for HET, and 92%, 4%, 1% for KO. **B.** Line graphs show mean square root residual (MSRR) calculated for the number of NMF components. The number of significant NMF components (dashed gray vertical line) was determined when the MSRR for the observed activity (blue line) in A was greater than the MSRR for randomized activity (gray solid line). **C-D.** Box plots show median (horizontal line), interquartile range (box), and range excluding outliers (vertical lines) with scatter plots (colored circles for each culture), and density curves for the number of significant NMF components (**C**) and mean number of electrodes detecting activity from neurons in each significant NMF component (**D**). * p≤0.05, ** p≤0.01 (nparLD, Dunn’s Test).

## Discussion

### Main findings

We show that Mecp2 deficiency alters the developmental trajectory of in vitro microscale murine cortical networks. The developmental increase in firing and network bursts rates, functional connectivity, network size, and density is delayed and diminished in the cultures heterozygous and hemizygous for the *Mecp2* deletion compared to the wildtype cultures. This has a significant effect on network metrics for local and global information processing, even when normalized for differences in network size and density using surrogate networks for comparison. These altered network metrics predict impaired capacity for cellular-scale information, which we show evidence from using non-negative matrix factorization to identify and quantify the relative influence on patterns of activity from subnetworks in the overall network activity. These network features, taken together, reveal microscale impairment in the development of networks, and subnetworks, that can support efficient and multiple streams of information processing.

### Mecp2 deficiency alters functional connectivity at multiple spatial scales

The altered trajectories of network development at the microscale may underlie network-level deficits at larger spatial scales. Imaging studies of functional networks, using optical imaging of the whole cortex, revealed differences in the overall functional connectivity and node degree in Mecp2-deficient mice [18]. Anatomical studies revealed regional differences in the development of long-range connectivity, as seen in anatomical studies of the developmental hippocampal circuits in mice [19]. Whole-brain or regional differences in network architecture have also been observed in EEG recordings from children with Rett syndrome [5]. Our in vitro microscale approach reveals how the trajectories of network development diverge early in brain development. This is particularly relevant for the timing of therapeutic interventions. These studies predict that gene therapies to restore MeCP2 function would need to be applied very early in development to ensure networks that can support complex information processing. In contrast, for children and adults in which the brain networks have already formed, increasing MeCP2 expression can improve neuronal maintenance functions but additional downstream therapeutic strategies will be necessary to modulate network function.

### Mosaic cultures reveal non-cell-autonomous effects of Mecp2 deficiency

Due to X-inactivation in females with Rett syndrome, *MECP2* heterozygosity results in cellular mosaicism with each cell expressing either the normal or mutant MeCP2 protein, rather than each cell expressing 50% of the normal level of MeCP2. Thus, in our murine cultures, we were able to compare the effect on network development of having no cells (Mecp2-KO), a mix of cells (Mecp2-Het), and all cells (Mecp2-WT) expressing Mecp2. One of the key questions in elucidating the mechanisms underlying cortical information-processing deficits in Rett syndrome, and developing strategies to improve function, is whether the effects of MeCP2 deficiency are cell-autonomous (affecting only those cells without functional MeCP2) or non-cell-autonomous (affecting those cells in the network expressing functional MeCP2 as well). Our findings reveal many network features that were similarly impaired in the Mecp2-Het and -KO cultures, supporting non-cell-autonomous effects of Mecp2 deficiency in the development of murine cortical circuits. Moreover, the Mecp2-Het networks also diverge from the Mecp2-KO, for example in the developmental trajectory of mean network burst rates and rich-club topology. Non-cell-autonomous effects of Mecp2 deficiency have also been observed at the transcriptomic level in the hippocampus [24]. Neuronal networks develop in culture, driven by their genetic programming, to support complex spatiotemporal information processing [20]. Our findings show that mosaic expression of a Mecp2 loss-of-function mutation is sufficient to alter how functional microscale networks form and their ability to support information sharing. Thus, correcting MeCP2 expression levels in human cells expressing the mutant *MECP2* postnatally, for example, may not be sufficient to rescue cortical processing deficits. These non-cell-autonomous effects also support the need for strategies that modulate network function, rather than gene expression alone, to restore cortical function in children and adults with Rett syndrome.

### Developmental trajectory key for elucidating network deficits

Our study compares the developmental trajectory of neuronal activity and microscale network function from early postnatal murine cultures over the first 5 weeks in vitro. This work reveals that, although there are delays in multiple features in the Mecp2-deficient networks, these networks do mature. The topological differences, including the smaller rich clubs, predict lower efficiency in both the Mecp2-Het and -KO networks compared to the WT. Interestingly, in an optical imaging study of whole-brain or regional-level network function in adult mice, there were opposing effects on node degree in male (hemizygous for *Mecp2* deletion) and female (heterozygous for *Mecp2* deletion) [18]. This difference is likely explained by the later symptom onset in female mice. Our study represents early postnatal development, in which we can already detect differences in the microscale networks that develop before the behavioral decline would occur in vivo. In the imaging study of juvenile and adult mice, Mecp2-deficient male mice typically show a behavioral decline between 5-8 weeks, whereas this same decline may occur 3-12 months later, depending on the X-inactivation skew, in the female mice. Thus, the sex-related differences in node degree observed at larger spatial scales [18] may be capturing different stages in the disease progression.

Differences between studies of neuronal activity, as measured by mean firing and burst rates, are also dependent on the developmental time points studied. Mecp2-KO murine neuronal precursor cells (NPCs) plated on MEAs at embryonic day (E)15 were assessed at DIV18 and 22 did not show the increase in number of active electrodes, mean firing or burst rates, as seen in the Mecp2-WT cultures [25]. Our study in early postnatal cortical cultures reveals that this is a delay in the developmental increase and that the low number of active electrodes and activity in those electrodes does increase with age. Electrophysiological studies of neuronal activity in acute brain slices or in vivo recordings from juvenile and adult Mecp2-deficient mice also are dependent on the developmental age, brain region, and whether the *MECP2* mutation led to a complete KO or was isolated to specific cell-types or developmental stages [26]. In this study, we used neonatal cortical cultures from mice with a loss-of-function *MECP2* mutation (deletion of exon 3 and 4). In vivo single-unit recordings in visual cortex in this mouse model showed a marked decrease in spontaneous and sensory-evoked activity in Mecp2-KO compared to WT littermates [14]. Early in vitro recordings from excitatory cells in somatosensory cortex also showed decreased spontaneous activity (2-fold less at postnatal day 14, 4-fold less after symptoms onset at day 28-35) in Mecp2-KO compared to WT in acute slices [7]. Cultured hippocampal neurons at DIV 14 also showed decreased spontaneous activity in Mecp2-KO compared to WT networks [10].

Studies of the effects of MeCP2 deficiency on human iPSC-derived cortical tissues also show differences in firing and burst rates that are also likely specific to the developmental timepoints assessed, which represent periods in human fetal brain development [27]. *MECP2*-null iPSC-derived cortical excitatory neuronal cultures in which overexpression of neurogenin 2 (NGN2) has been used to accelerate the development, skipping the neuronal precursor stages, show iPSC-line- and age-dependent effects on firing and burst rates in recordings between 3-7 weeks in culture [28], while other 2D *MECP2* knock-out models showed very little spiking compared to the isogenic control cultures at DIV 51-55 [29]. MEA recordings from *MECP2* knock-out human cerebral organoids showed reduced spontaneous activity at DIV30, as measured by population spiking, compared to controls [30]. Notably, neuronal networks in human cerebral organoids develop much more slowly than in murine cultures (over months in human-derived versus days-to-weeks in murine cultures). Thus, DIV30 in a human organoid is very early in embryonic development. In our own work with MEA recordings from control human cerebral organoids [31], we see more mature network activity starting around DIV180, for reference. Early development of hypersynchronous activity was observed with calcium imaging in MeCP2-deficient human cerebral organoids at DIV 70-100 [32], evidence of an altered trajectory of network development.

### Limitations of study

The differences in network activity, topology, and dynamics identified in this study were identified in MEA recordings from primary murine cultures. While they offer insight into the microscale functional networks in early postnatal development that is currently not possible to do in vivo, further studies will be necessary to see which features are also observed in vivo. This study provides an approach that can be applied to human-derived cultures; however, developmental time points from later stages of human cerebral organoid development (e.g., DIV 150-300) will be needed to see if similar developmental effects occur in human-derived cultures. In this study, we consider a node in the network to be the activity observed from a neuron, or neurons, near an individual electrode, without spike sorting. Network topological features can be seen across spatial scales in the brain; however, future studies could combine simultaneous calcium imaging with MEA recording to provide ground truth for the number of active units near individual electrodes and to deconvolve the multi-unit to single-unit activity. Notably, a recent study shows that network dynamics can be accurately estimated from multi-unit activity without spike sorting [33].

The inference of functional connectivity was made using the spike time tiling coefficient [22], which was designed specifically for MEA recordings and is largely rate independent in estimating correlation values. Spike timing underlies synaptic communication and plasticity in the brain. Thus, MEA recordings have the advantage over calcium imaging in high sensitivity and precision of spike timing (in milliseconds) for inferring functional connectivity. Calcium imaging provides single-cell resolution, but relies on a transient rise in fluorescence triggered by movement of calcium as a proxy for cell activity with a much lower temporal resolution (typically seconds). Thus, calcium imaging would likely be able to detect differences in burst rates, but not differences in action potential firing between bursts that would impact the inference of functional connectivity with MEA recordings, but not with calcium imaging. The dimensional reduction approach, using non-negative matrix factorization, does not rely on pair-wise correlations to infer functional connectivity. Thus, it provides a complementary approach to comparing network topology using graph theoretical metrics. The reduction in the number of activity patterns (significant NMF components) in the Mecp2-deficient networks provides additional evidence of impairments in network function, which were predicted by the topological differences in the developing microscale networks.

### Broader application

The network topology and dynamics metrics we applied to microscale murine Mecp2-deficient and wild-type cortical circuits are commonly used in network science to examine the efficiency and information-sharing capacity of biological, and non-biological, networks at different spatial scales. However, they have historically been underutilized for microscale networks including MEA recordings [16]. The altered trajectory of network development identified here provides mechanistic insight into impairments in cortical processing that have been observed at higher scale scales (e.g., EEG) and are apparent in individuals with Rett syndrome, particularly as the disorder progresses. These differences in network topology and dynamics in MEA recordings from murine, or human-derived neuronal cultures, can also provide a platform for testing new therapeutic products for restoring network function in Rett syndrome.

## Methods

### RESOURCE AVAILABILITY

#### Lead contact

Further information and requests for resources should be directed to the lead contact, Susanna Mierau.

## Materials availability

This study did not generate new unique reagents.

## Data and code availability

### Data

The MEA spike time data has been deposited at the Harvard Dataverse (https://doi.org/10.7910/DVN/FVCM4S) and is publicly available.

### Code

All original code for analysis and production of figures in this paper have been deposited in the Harvard Dataverse (https://doi.org/10.7910/DVN/FVCM4S) and are publicly available. The latest version of MEA-NAP is publicly available at GitHub (https://github.com/SAND-Lab/MEA-NAP).

## EXPERIMENTAL MODEL

### Murine 2D cultures

Female Mecp2-heterozygous mice (Jackson Laboratory, Strain 003890) were bred in house, with approval from the UK Home Office, with either C57BL/6J or double homozygote for PV-Cre (B6;129P2-Pvalb^tm1(cre)Arbr^/J, Jackson Laboratory, Strain 008069) and tdTomato-flox (B6.Cg-*Gt(ROSA)26Sor^tm14(CAG-tdTomato)Hze^*/J, Strain 007914) males. Offspring were either wild-type, heterozygous, hemizygous for Mecp2 deletion. Male and female neonatal mice were sacrificed at postnatal day 0 in accordance with UK Home Office regulations by first inducing hypothermia prior to decapitation. The cortices were dissected in sterile ice-cold phosphate buffer solution (PBS, Gibco, 14190094) under a stereoscope and were next chemically dissociated using a 1:1 mixture of papain (Sigma, P5306) and sterile PBS at 37 °C for 25 minutes. The reaction was stopped by adding neurobasal-A media (Gibco, 10888022) with B27 supplement (NB-B27; B27 supplement, Gibco, 17504044) with 4% bovine serum albumin (BSA, Sigma, A8412), and the cortices were manually dissociated using a pipette. A small volume (20 μL) of the cells were removed for manual cell counting with a hemocytometer, and the remaining volume was then centrifuged for 10 minutes. Before the supernatant was removed, DNase I (Roche, 10104159001) was added for 5 minutes to reduce cell aggregation. The pellet was resuspended in neurobasal-A media with the B27 supplement (NB-B27) and 0.25% Glutamax (Gibco, 35050-038). The cells were resuspended in the volume necessary to ensure that 30 μL would contain 5x10^4^ cells, which were plated on single-well 60-channel MEA chips (Multi-channel systems, 60MEA200/30iR-ITO-gr).

### MEA preparation and plating

The MEA chips were treated in advance with heavy-weight poly-L-lysine (PLL; Sigma, P4832) for 5 minutes up to 24 hours at 37 °C in the incubator, followed by three PBS washes to ensure full removal of the PLL, before adding 5 μL of laminin (Sigma, L2020) directly over the MEA grid. A 30 μL aliquot of 5x10^4^ cells was added directly to the laminin on the MEA grid, and the MEA chips were incubated for 30 minutes. Visual inspection under a light microscope was used to confirm cell adhesion prior to adding 565 μL of NB-B27 with 0.25% Glutamax at 37 °C to the MEA well. Cultures were maintained in the incubator at 37 °C with humidity control and 5% carbon dioxide. One-third of the media (180-200 μL/well) was exchanged three times per week with fresh NB-B27 with 0.25% Glutamax at 37 °C. Cells were visualized each week under the microscope prior to recording. Cultures with less than 75% of the electrodes covered by cells were discarded.

### MEA data acquisition

MEA recordings were made weekly from DIV 7-35 using the MEA2100 system (Multi-channel systems). We acquired MEA data using an MEA2100 dual headstage MEA system (Multi-channel systems), with the temperature controller (Multi-channel systems, TCX-2) set to 37 °C. Raw data was acquired at 25 kHz for 10 minutes using the MCRack software (Multi-channel systems). This duration of MEA recordings has been sufficient to reveal network development in prior studies [23,34]. Raw data was exported using MCTool (Multi-channel systems) and converted to MATLAB format (.mat) using custom scripts included in our MEA network analysis pipeline (MEA-NAP) [23].

### MEA data analysis

MEA data analysis was performed using custom scripts or MEA-NAP [23]. The voltage time series were first bandpass filtered (third-order Butterworth filter, 600-8000 Hz). Template-based spike detection was performed using the bior1.5 template in MATLAB with a cost parameter of -0.0627. This cost parameter was selected by comparing spike detection before and after treatment of cultures with tetrodotoxin (TTX, n=6). The mean firing rate for each electrode was calculated as the number of action potentials divided by the total length of the recording (in seconds), which in turn was averaged for each recording. The number of active channels was calculated as the number of electrodes per MEA in which the number of spikes detected divided by the length of the recording was greater than 0.01 Hz. Bursts within individual electrodes were detected using the ISI_N_ method [35], with the threshold set automatically based on the interspike interval distribution. Network bursts were defined as a minimum of 10 spikes in at least 3 channels, and the ISI_N_ threshold set automatically [35].

Functional connectivity was inferred using the STTC with a 10 ms coincidence window. To determine significant pairwise connections, probabilistic thresholding was performed with the threshold for significance where the real weight was ≤0.1% of the distribution of edge weights from circular shifts (n=2000). Graph theoretical metrics were calculated as in MEA-NAP [22] with the exceptions of betweenness centrality and small-worldness coefficient, in which surrogate graphs (n=200 per network) were used to normalize the metrics. Surrogate graphs were created by taking the observed graph and rewiring edges while preserving the node degree distribution using the Maslov-Sneppen algorithm in the Brain Connectivity Toolbox [36]. The rich-club coefficient was calculated as the proportion of edges realized among rich-club nodes [34]. Rich-club nodes were defined as a node with greater than equal to k edges. We selected the k value that gave the highest rich-club coefficient across all values of k. The maximum value of k is the maximum node degree in the network. This was also the k value used for the corresponding surrogate graphs to normalize the rich-club coefficient.

Non-negative matrix factorization (NMF) was performed on the spike train data down-sampled to 10 Hz using MEA-NAP [23]. The number of significant NMF components was determined by first calculating the mean square root residual (MSRR) between the observed activity and the rank-k approximation. We then shuffled one the activity of each electrode across time bins to destroy temporal correlations between channels to create a random network and calculated the MSRR of the rank-k approximation. We then identified the maximum number of significant NMF components for which the MSRR in the observed activity was greater than in the randomized network. For each NMF component, the number of active electrodes (mean firing rate greater than 0.01 Hz) was used to calculate the mean size of the subnetworks for the significant NMF components in the observed network activity.

### Data visualization

Figure panels were created in MATLAB using custom scripts (A.W.E.D.) and MEA-NAP [23].

### Immunohistochemistry

Sterile coverslips pre-treated with poly-d-lysine and laminin (12 mm diameter, Neuvitro Corp., GG-12-1.5-Laminin) were plated with 1.5 x 10^5^ cells and maintained under similar conditions in a 24-well plate (Corning, 3524) to the cultures plated on the MEA. Cells were fixed by first aspirating the media and then adding 300 μL of a mixture of warm (37 °C) 4% paraformaldehyde (PFA, Thermo Fisher Scientific, J61899-AK) and 4% sucrose (Fisher Scientific, 10634932, S/8600/60) in PBS. The coverslip was incubated in the PFA/sucrose mixture at room temperature for 10 minutes and then replaced with 300 μL of 50 mM ammonium chloride (NH_4_Cl, Thermo Fisher Scientific, A15000) for 10 minutes. Coverslips were then washed with DPBS (Gibco, 14190-094) and treated with 300 μL of 0.1% Triton X-100 (Sigma-Aldrich, T9284) in DPBS for 10 minutes. Next coverslips were washed with DPBS and incubated for 30 minutes with 300 μL of 4% BSA (Appleton Woods, CSR602) in DPBS on a shaker. Primary and secondary antibodies for Mecp2 (Rat anti-Mecp2, clone 4H7, Millipore Sigma, MABE328; Alexa Fluor goat anti-rat 568, Invitrogen, A-11077), MAP2 (Chicken anti-MAP2, Millipore Sigma, AB5543; Goat anti-chicken IgY Alexa Fluor 488, Thermo Fisher Scientific, A-11039), and DAPI (Fluoroshield with DAPI, Scientific Laboratory Supplies, F6057) were used. 600 μL of the primary antibody was diluted in 10% BSA/PBS mixture and incubated overnight at 4 °C for overnight on a shaker. Coverslips were washed thrice for 5 minutes each with DPBS on a shaker at room temperature. Fluorescently conjugated secondary antibodies were also diluted in 10% BSA/DPBS and 600 μLadded to each coverslip followed by a 1-hour incubation at room temperature on the shaker. After 3 more DPBS washes, the coverslips were mounted with cover glass using Fluoroshield antifade mounting medium without DAPI (Sigma-Aldrich, F6182).

## QUANTIFICATION AND STATISTICAL ANALYSIS

Statistical tests were completed in R. Data were tested for normality using the Shapiro-Wilk test [37] and through visual inspection of Q–Q plots. After fitting a linear model to the data in mixed design experiments, the normality assumption was checked based on the residuals of this model and the sphericity assumption using Mauchly’s test [38]. Where these assumptions were violated, a non-parametric approach was employed [39,40]. As assumptions for electrophysiological and graph metrics were not always met, a non-parametric approach was used across all metrics for consistency. This included a non-parametric ANOVA-type statistic (nparLD) equivalent to a mixed ANOVA calculated on ranks rather than raw data [41]. Significant age-genotype interactions and significant main effects of genotype were followed up with post-hoc comparisons between pairs of groups at each timepoint were made with Dunn’s Test [42]. This controls the family-wise error rate (FWER) for multiple comparisons and is typically used with nonparametric multi-group analyses such as the Kruskal-Wallis test [42,43].

## ACKNOWLEDGMENTS

We would like to thank our funders: the European Commission Horizon 2020 (Marie Skłodowska-Curie Actions Individual European Fellowship, EU Project 700999, S.B.M.), the American Academy of Neurology (Career Development Award, S.B.M.), and the UKRI Medical Research Council (Doctoral Training Partnership, A.W.E.D.). S.B.M. is supported by an NIH NINDS K02 Independent Scientist Award (1K02NS131521-01A1). Thank you to animal welfare and husbandry staff at the University of Cambridge and Wellcome Sanger Institute.

## CONTRIBUTIONS

Concept: S.B.M, O.P., A.W.E.D., T.P.H.S.

Methodology: A.W.E.D., T.P.H.S., R.C.F., A.S., R.T., S.J.E., O.P., S.B.M.

Investigation: A.W.E.D., R.C.F., Y.Y., A.S., I.L., S.B.M.

Formal analysis: A.W.E.D., T.P.H.S., S.B.M.

Visualization: A.W.E.D., T.P.H.S., R.C.F., S.B.M.

Writing – original draft: S.B.M., A.W.E.D.

Writing – reviewing & editing: A.W.E.D., T.P.H.S., S.B.M., S.J.E., O.P. Funding acquisition: S.B.M., O.P., A.W.E.D.

Supervision: S.B.M., S.J.E., O.P.

---

**Table S1.** Statistical comparisons of neuronal activity in Mecp2-deficient and wildtype microscale networks, Related to. **Figure 1**. Number of active electrodes and mean firing rate in Mecp2-WT, -Het, and -KO cortical cultures (n=number of cultures) from DIV 14–35. Array firing rate is defined as the mean across recordings of the median electrode firing rate (spikes/s) per recording. Dunn’s test post hoc pairwise comparisons between genotypes are shown for each DIV following nparLD analysis in addition to Cohen’s *d* effect size statistic for each comparison.

| Summary |  | # Active Electrodes |  |  | Array Firing Rate |  |  |
| --- | --- | --- | --- | --- | --- | --- | --- |
| Group | DIV | Mean | ±SEM |  | Mean | ±SEM |  |
| WT<br>n=18 | 14 | 40.778 | 1.933 |  | 2.944 | 0.560 |  |
|  | 21 | 48.500 | 1.522 |  | 5.481 | 1.067 |  |
|  | 28 | 51.500 | 1.153 |  | 10.120 | 1.938 |  |
|  | 35 | 49.722 | 2.066 |  | 11.107 | 1.783 |  |
| HET<br>n=19 | 14 | 31.053 | 2.937 |  | 1.033 | 0.177 |  |
|  | 21 | 41.579 | 2.432 |  | 2.734 | 0.627 |  |
|  | 28 | 43.053 | 2.846 |  | 5.021 | 1.059 |  |
|  | 35 | 39.000 | 3.891 |  | 5.289 | 1.524 |  |
| KO<br>n=15 | 14 | 30.200 | 3.509 |  | 1.144 | 0.279 |  |
|  | 21 | 39.533 | 3.681 |  | 3.168 | 1.055 |  |
|  | 28 | 43.733 | 3.454 |  | 6.748 | 2.346 |  |
|  | 35 | 38.933 | 4.726 |  | 6.395 | 1.894 |  |
| Dunn's test *p* |  | WT-HET | WT-KO | HET-KO | WT-HET | WT-KO | HET-KO |
|  | 14 | 0.003 | 0.004 | 0.492 | 0.009 | 0.007 | 0.405 |
|  | 21 | 0.052 | 0.019 | 0.292 | 0.033 | 0.035 | 0.468 |
|  | 28 | 0.037 | 0.020 | 0.351 | 0.015 | 0.042 | 0.372 |
|  | 35 | 0.011 | 0.019 | 0.466 | 0.012 | 0.045 | 0.335 |
| Cohen's *d* |  |  |  |  |  |  |  |
|  | 14 | 0.899 | 0.964 | 0.065 | 1.094 | 0.945 | 0.120 |
|  | 21 | 0.783 | 0.837 | 0.166 | 0.740 | 0.534 | 0.128 |
|  | 28 | 0.887 | 0.801 | 0.053 | 0.770 | 0.391 | 0.249 |
|  | 35 | 0.788 | 0.776 | 0.004 | 0.819 | 0.632 | 0.159 |
| Legend: days in vitro (DIV); SEM (standard error of the mean); wildtype (WT); Mecp2-heterozygous (HET); Mecp2-hemizygous (KO) |  |  |  |  |  |  |  |

**Table S2.**
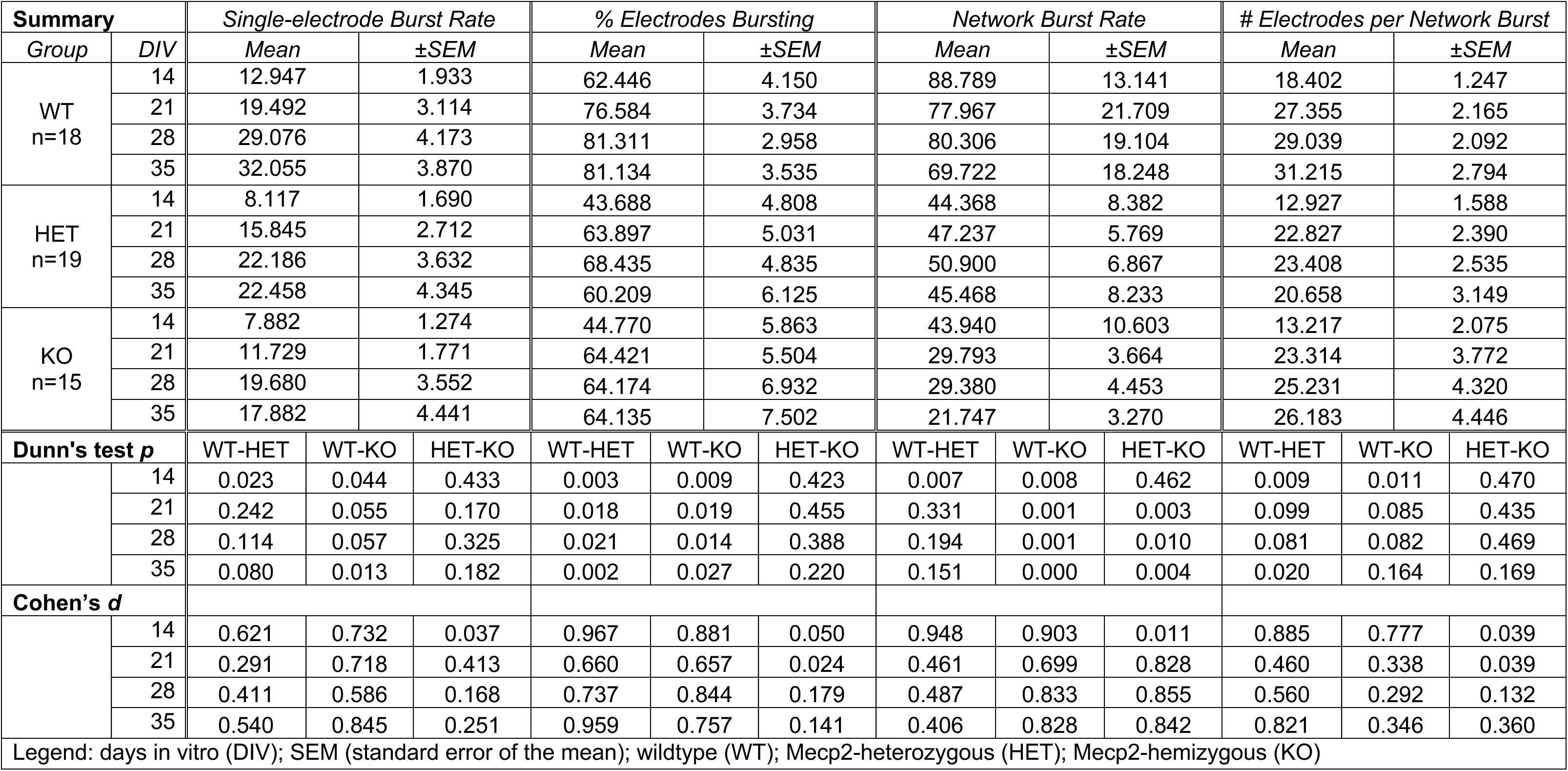
Statistical comparisons of bursting in Mecp2-deficient and wildtype microscale networks, Related to. **Figure 2**. Single-electrode bursting and network burst metrics in Mecp2-WT, -Het, and -KO cortical cultures (n=number of cultures) across DIV 14-35. Reported measures include single-electrode burst rate, percentage of bursting electrodes, network burst rate, and number of electrodes recruited per burst. Dunn’s test post hoc pairwise comparisons between genotypes are shown for each DIV following nparLD analysis, along with Cohen’s d effect size statistics.

| Summary |  | Single-electrode Burst Rate |  |  | % Electrodes Bursting |  |  | Network Burst Rate |  |  | # Electrodes per Network Burst |  |  |
| --- | --- | --- | --- | --- | --- | --- | --- | --- | --- | --- | --- | --- | --- |
| Group | DIV | Mean | ±SEM |  | Mean | ±SEM |  | Mean | ±SEM |  | Mean | ±SEM |  |
| WT<br>n=18 | 14 | 12.947 | 1.933 |  | 62.446 | 4.150 |  | 88.789 | 13.141 |  | 18.402 | 1.247 |  |
|  | 21 | 19.492 | 3.114 |  | 76.584 | 3.734 |  | 77.967 | 21.709 |  | 27.355 | 2.165 |  |
|  | 28 | 29.076 | 4.173 |  | 81.311 | 2.958 |  | 80.306 | 19.104 |  | 29.039 | 2.092 |  |
|  | 35 | 32.055 | 3.870 |  | 81.134 | 3.535 |  | 69.722 | 18.248 |  | 31.215 | 2.794 |  |
| HET<br>n=19 | 14 | 8.117 | 1.690 |  | 43.688 | 4.808 |  | 44.368 | 8.382 |  | 12.927 | 1.588 |  |
|  | 21 | 15.845 | 2.712 |  | 63.897 | 5.031 |  | 47.237 | 5.769 |  | 22.827 | 2.390 |  |
|  | 28 | 22.186 | 3.632 |  | 68.435 | 4.835 |  | 50.900 | 6.867 |  | 23.408 | 2.535 |  |
|  | 35 | 22.458 | 4.345 |  | 60.209 | 6.125 |  | 45.468 | 8.233 |  | 20.658 | 3.149 |  |
| KO<br>n=15 | 14 | 7.882 | 1.274 |  | 44.770 | 5.863 |  | 43.940 | 10.603 |  | 13.217 | 2.075 |  |
|  | 21 | 11.729 | 1.771 |  | 64.421 | 5.504 |  | 29.793 | 3.664 |  | 23.314 | 3.772 |  |
|  | 28 | 19.680 | 3.552 |  | 64.174 | 6.932 |  | 29.380 | 4.453 |  | 25.231 | 4.320 |  |
|  | 35 | 17.882 | 4.441 |  | 64.135 | 7.502 |  | 21.747 | 3.270 |  | 26.183 | 4.446 |  |
| Dunn's test *p* |  | WT-HET | WT-KO | HET-KO | WT-HET | WT-KO | HET-KO | WT-HET | WT-KO | HET-KO | WT-HET | WT-KO | HET-KO |
|  | 14 | 0.023 | 0.044 | 0.433 | 0.003 | 0.009 | 0.423 | 0.007 | 0.008 | 0.462 | 0.009 | 0.011 | 0.470 |
|  | 21 | 0.242 | 0.055 | 0.170 | 0.018 | 0.019 | 0.455 | 0.331 | 0.001 | 0.003 | 0.099 | 0.085 | 0.435 |
|  | 28 | 0.114 | 0.057 | 0.325 | 0.021 | 0.014 | 0.388 | 0.194 | 0.001 | 0.010 | 0.081 | 0.082 | 0.469 |
|  | 35 | 0.080 | 0.013 | 0.182 | 0.002 | 0.027 | 0.220 | 0.151 | 0.000 | 0.004 | 0.020 | 0.164 | 0.169 |
| Cohen's *d* |  |  |  |  |  |  |  |  |  |  |  |  |  |
|  | 14 | 0.621 | 0.732 | 0.037 | 0.967 | 0.881 | 0.050 | 0.948 | 0.903 | 0.011 | 0.885 | 0.777 | 0.039 |
|  | 21 | 0.291 | 0.718 | 0.413 | 0.660 | 0.657 | 0.024 | 0.461 | 0.699 | 0.828 | 0.460 | 0.338 | 0.039 |
|  | 28 | 0.411 | 0.586 | 0.168 | 0.737 | 0.844 | 0.179 | 0.487 | 0.833 | 0.855 | 0.560 | 0.292 | 0.132 |
|  | 35 | 0.540 | 0.845 | 0.251 | 0.959 | 0.757 | 0.141 | 0.406 | 0.828 | 0.842 | 0.821 | 0.346 | 0.360 |
| Legend: days in vitro (DIV); SEM (standard error of the mean); wildtype (WT); Mecp2-heterozygous (HET); Mecp2-hemizygous (KO) |  |  |  |  |  |  |  |  |  |  |  |  |  |

**Table S3.** Statistical comparisons of network features in Mecp2-deficient and wildtype microscale networks, Related to. **Figures 3**. Graph-theoretic properties of functional networks derived from MEA recordings in Mecp2-WT, -Het, and -KO cortical cultures (n=number of cultures) across DIV 14-35. Metrics include mean edge weight, mean node degree, network size, and network density. Statistical comparisons between genotypes were performed using nparLD with Dunn’s test post hoc comparisons and Cohen’s d effect sizes.

| Summary |  | Mean Edge Weight |  |  | Mean Node Degree |  |  | Network Size |  |  | Network Density |  |  |
| --- | --- | --- | --- | --- | --- | --- | --- | --- | --- | --- | --- | --- | --- |
| Group | DIV | Mean | ±SEM |  | Mean | ±SEM |  | Mean | ±SEM |  | Mean | ±SEM |  |
| WT<br>n=18 | 14 | 0.202 | 0.023 |  | 32.822 | 2.884 |  | 40.444 | 2.000 |  | 0.805 | 0.040 |  |
|  | 21 | 0.415 | 0.028 |  | 45.312 | 1.976 |  | 48.333 | 1.510 |  | 0.950 | 0.016 |  |
|  | 28 | 0.446 | 0.029 |  | 48.216 | 1.713 |  | 51.333 | 1.149 |  | 0.954 | 0.018 |  |
|  | 35 | 0.454 | 0.035 |  | 46.392 | 2.716 |  | 49.611 | 2.068 |  | 0.937 | 0.027 |  |
| HET<br>n=19 | 14 | 0.125 | 0.021 |  | 20.967 | 3.274 |  | 29.842 | 3.212 |  | 0.651 | 0.045 |  |
|  | 21 | 0.297 | 0.040 |  | 34.586 | 3.417 |  | 40.947 | 2.648 |  | 0.823 | 0.045 |  |
|  | 28 | 0.307 | 0.040 |  | 36.223 | 3.518 |  | 42.474 | 3.024 |  | 0.833 | 0.039 |  |
|  | 35 | 0.272 | 0.045 |  | 32.075 | 4.366 |  | 38.053 | 4.151 |  | 0.751 | 0.070 |  |
| KO<br>n=15 | 14 | 0.136 | 0.031 |  | 20.269 | 4.233 |  | 28.867 | 3.942 |  | 0.597 | 0.072 |  |
|  | 21 | 0.299 | 0.047 |  | 33.474 | 4.325 |  | 39.267 | 3.760 |  | 0.814 | 0.049 |  |
|  | 28 | 0.330 | 0.059 |  | 35.043 | 4.494 |  | 43.267 | 3.592 |  | 0.778 | 0.062 |  |
|  | 35 | 0.354 | 0.061 |  | 34.707 | 5.271 |  | 37.800 | 5.032 |  | 0.913 | 0.042 |  |
| Dunn's test *p* |  | WT-HET | WT-KO | HET-KO | WT-HET | WT-KO | HET-KO | WT-HET | WT-KO | HET-KO | WT-HET | WT-KO | HET-KO |
|  | 14 | 0.014 | 0.022 | 0.482 | 0.009 | 0.007 | 0.414 | 0.009 | 0.007 | 0.394 | 0.013 | 0.008 | 0.386 |
|  | 21 | 0.015 | 0.022 | 0.488 | 0.013 | 0.019 | 0.492 | 0.035 | 0.038 | 0.474 | 0.010 | 0.005 | 0.343 |
|  | 28 | 0.010 | 0.045 | 0.306 | 0.010 | 0.015 | 0.486 | 0.015 | 0.045 | 0.361 | 0.005 | 0.004 | 0.390 |
|  | 35 | 0.003 | 0.108 | 0.092 | 0.006 | 0.040 | 0.262 | 0.010 | 0.033 | 0.368 | 0.002 | 0.494 | 0.003 |
| Cohen's *d* |  |  |  |  |  |  |  |  |  |  |  |  |  |
|  | 14 | 0.816 | 0.609 | 0.106 | 0.890 | 0.880 | 0.046 | 0.910 | 0.962 | 0.067 | 0.839 | 0.920 | 0.231 |
|  | 21 | 0.787 | 0.779 | 0.007 | 0.881 | 0.921 | 0.071 | 0.785 | 0.834 | 0.130 | 0.865 | 0.988 | 0.044 |
|  | 28 | 0.919 | 0.651 | 0.116 | 0.991 | 1.023 | 0.073 | 0.882 | 0.805 | 0.059 | 0.905 | 1.034 | 0.270 |
|  | 35 | 1.049 | 0.518 | 0.384 | 0.904 | 0.723 | 0.134 | 0.806 | 0.808 | 0.013 | 0.799 | 0.176 | 0.641 |
| Legend: days in vitro (DIV); SEM (standard error of the mean); wildtype (WT); Mecp2-heterozygous (HET); Mecp2-hemizygous (KO) |  |  |  |  |  |  |  |  |  |  |  |  |  |

**Table S4.**
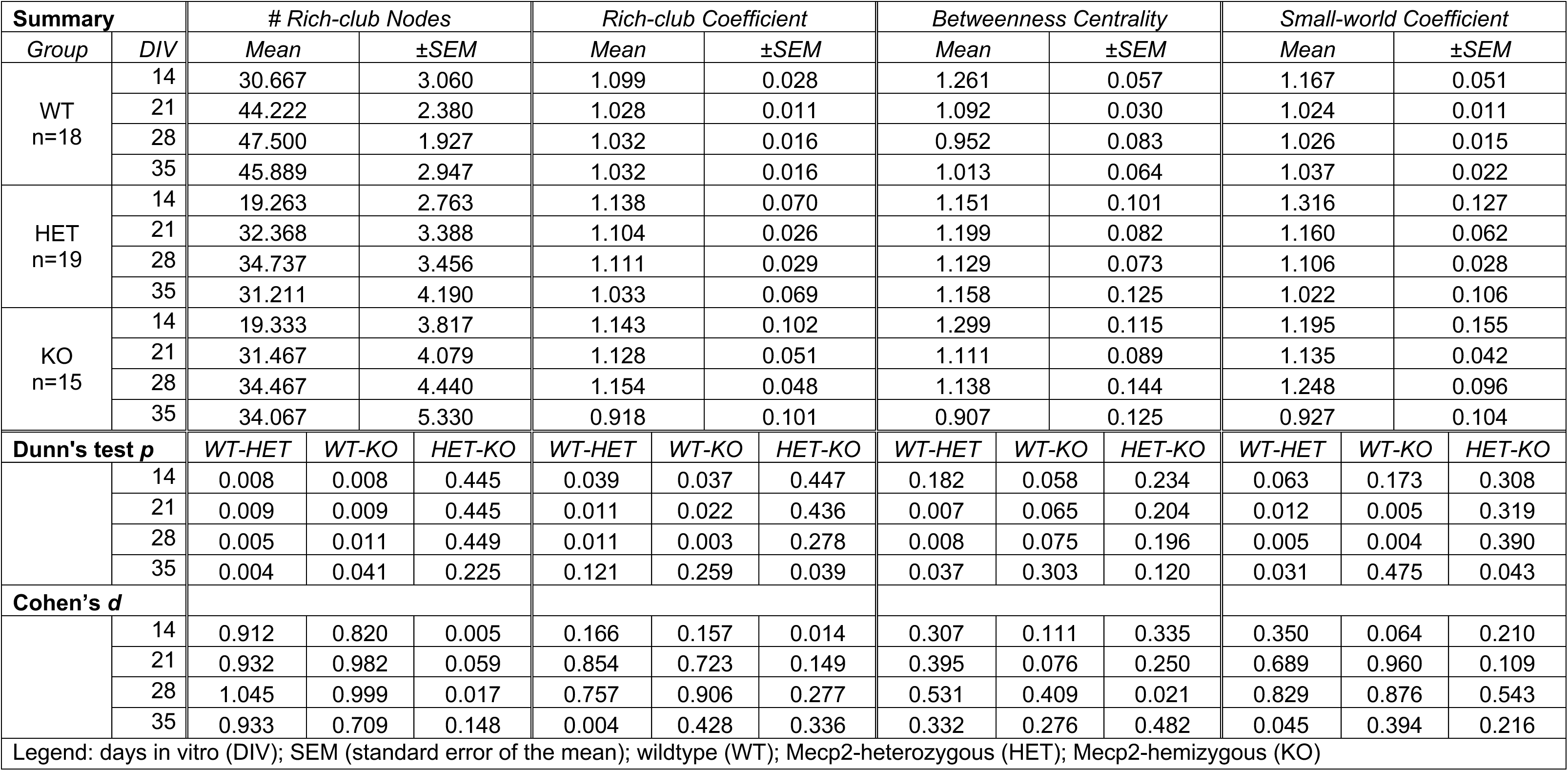
Statistical comparisons of rich-club hubs and small-world topology in Mecp2-deficient and wildtype microscale networks, Related to. **Figure 4**. Higher-order network organization metrics in Mecp2-WT, -Het, and -KO cortical cultures (n=number of cultures) across DIV 14-35. Measures include the number of rich-club nodes, rich-club coefficient, betweenness centrality, and small-world coefficient. Dunn’s test post hoc pairwise comparisons following nparLD analysis and Cohen’s d effect sizes are reported for each DIV.

**Table S5.** Statistical comparisons of network dynamics in Mecp2-deficient and wildtype microscale networks, Related to. **Figure 5**. Properties of activity patterns identified using non-negative matrix factorization (NMF) in Mecp2-WT, -Het, and -KO cortical cultures (n=number of cultures) across DIV 14-35. Reported measures include the number of significant NMF components and the number of electrodes per significant NMF component. Genotype comparisons were assessed using nparLD with Dunn’s test post hoc comparisons and Cohen’s d effect sizes.

| Summary |  | # Significant NMF Components |  |  | # Electrodes Per NMF Component |  |  |
| --- | --- | --- | --- | --- | --- | --- | --- |
| Group | DIV | Mean | ±SEM |  | Mean | ±SEM |  |
| WT<br>n=18 | 14 | 22.889 | 1.762 |  | 8.075 | 1.384 |  |
|  | 21 | 24.889 | 1.885 |  | 15.158 | 2.121 |  |
|  | 28 | 26.889 | 1.475 |  | 16.829 | 0.901 |  |
|  | 35 | 25.889 | 1.831 |  | 17.352 | 1.422 |  |
| HET<br>n=19 | 14 | 13.842 | 1.898 |  | 6.024 | 1.028 |  |
|  | 21 | 19.421 | 1.728 |  | 10.996 | 1.532 |  |
|  | 28 | 20.526 | 1.886 |  | 13.043 | 1.599 |  |
|  | 35 | 18.000 | 2.116 |  | 12.377 | 2.717 |  |
| KO<br>n=15 | 14 | 13.533 | 2.428 |  | 6.352 | 1.576 |  |
|  | 21 | 18.867 | 2.453 |  | 10.801 | 2.149 |  |
|  | 28 | 19.667 | 2.860 |  | 14.185 | 2.636 |  |
|  | 35 | 20.667 | 2.700 |  | 11.332 | 2.395 |  |
| Dunn's test *p* |  | WT-HET | WT-KO | HET-KO | WT-HET | WT-KO | HET-KO |
|  | 14 | 0.002 | 0.002 | 0.436 | 0.155 | 0.121 | 0.413 |
|  | 21 | 0.009 | 0.018 | 0.443 | 0.060 | 0.069 | 0.492 |
|  | 28 | 0.008 | 0.023 | 0.395 | 0.092 | 0.275 | 0.255 |
|  | 35 | 0.008 | 0.085 | 0.180 | 0.016 | 0.027 | 0.461 |
| Cohen's *d* |  |  |  |  |  |  |  |
|  | 14 | 1.146 | 1.114 | 0.035 | 0.394 | 0.288 | 0.062 |
|  | 21 | 0.704 | 0.691 | 0.066 | 0.527 | 0.501 | 0.026 |
|  | 28 | 0.868 | 0.824 | 0.090 | 0.669 | 0.356 | 0.134 |
|  | 35 | 0.923 | 0.575 | 0.273 | 0.525 | 0.785 | 0.097 |
| Legend: days in vitro (DIV); SEM (standard error of the mean); wildtype (WT); Mecp2-heterozygous (HET); Mecp2-hemizygous (KO); non-negative matrix factorization (NMF) |  |  |  |  |  |  |  |

